# Consensus genomic regions associated with multiple abiotic stress tolerance in wheat and implications for wheat breeding

**DOI:** 10.1101/2022.06.24.497482

**Authors:** Mohammad Jafar Tanin, Dinesh Kumar Saini, Karansher Singh Sandhu, Neeraj Pal, Santosh Gudi, Jyoti Chaudhary, Achla Sharma

**Author notes:** Corresponding author: Mohammad Jafar Tanin, Email ID. These authors contributed equally to this work (Mohammad Jafar Tanin, Dinesh Kumar Saini).

## Abstract

In wheat, a meta-analysis was performed using previously identified QTLs associated with drought stress, heat stress, salinity stress, water-logging stress, pre-harvest sprouting, and aluminium stress which predicted a total of 134 meta-QTLs (MQTLs) that involved at least 28 consistent and stable MQTLs conferring tolerance to five or all six abiotic stresses under study. Seventy-six MQTLs out of the 132 physically anchored MQTLs were also verified with genome-wide association studies. Around 43% of MQTLs had genetic and physical confidence intervals of less than 1 cM and 5 Mb, respectively. Consequently, 539 genes were identified in some selected MQTLs providing tolerance to 5 or all 6 abiotic stresses. Comparative analysis of genes underlying MQTLs with four RNA-seq based transcriptomic datasets unravelled a total of 191 differentially expressed genes which also included at least 11 most promising candidate genes common among different datasets. The promoter analysis showed that the promoters of these genes include many stress responsiveness cis-regulatory elements, such as ARE, MBS, TC-rich repeats, As-1 element, STRE, LTR, WRE3, and WUN-motif among others. Further, some MQTLs also overlapped with as many as 34 known abiotic stress tolerance genes. In addition, numerous ortho-MQTLs among the wheat, maize, and rice genomes were discovered. These findings could help with fine mapping and gene cloning, as well as marker-assisted breeding for multiple abiotic stress tolerances in wheat.

## Introduction

Continued crop improvement is paramount to feeding the continuously increasing human population, which is more critical regarding climate change, meeting sustainability goals, and limited natural resources^1,2^. Wheat is consumed by two-thirds of the world’s population, meeting 20% of dietary calories, and grown in a wide geographical distribution, 45 ^0^ S in Argentina to 67^0^ N in Scandinavia, including some high-altitude regions in the tropics and subtropics^3,4,5^. Owing to its wide distribution and climate variability, wheat is affected by various biotic (yellow, brown, and stem rusts, fusarium head blight, tan spot, and several other diseases, insects, and nematodes) and abiotic stresses (drought, heat, salinity, water-logging, pre-harvest sprouting, and mineral toxicity, among others)^6^. Breeding climate-resilient wheat cultivars is the best approach to assisting wheat in surviving abiotic stresses, which could be facilitated by mapping the genomic regions involved, marker-assisted breeding, and other advanced approaches such as genome editing, genomic-assisted breeding (involving the use of high-throughput genotyping and phenotyping systems), and haplotype-based breeding^7,8,4,5,9^.

Drought stress (DS), heat stress (HS), salinity stress (SS), water-logging stress (WS), pre-harvest sprouting (PHS), and aluminium stress (AS) are the major abiotic stresses affecting yield around the world^10,11^. Heat stress affects 58% of the wheat production area, while drought affects 42%^12^. Climatic uncertainty causes erratic rainfall, and warmer temperatures are projected in the future, which may convert long-season productive mega-environments into short-season heat stress environments^13^. These conditions, along with other abiotic stresses, represent a unique challenge to plant scientists in developing climatically resilient cultivars. Owing to the genetic (quantitative inheritance) and physiological complexity of tolerance traits, progress in enhancing tolerance or developing tolerant cultivars has been very slow. Furthermore, the wide variability of field conditions and the ineffectiveness of selection procedures impede progress. Traits contributing to tolerances to abiotic stresses are inherited quantitatively and controlled by a number of genetic loci. While numerous QTLs for each of the above-mentioned abiotic stress have been identified (http://www.wheatqtldb.net/), very few of the associated markers have been utilized in marker-assisted breeding (MAB) programmes owing to the comparatively lower heritability of the QTLs and other factors impacting the gene/QTL expression. Further, these studies are conditioned on different QTL mapping methods and various breeding methods, which makes it challenging to choose appropriate QTLs for breeding programmes that could confer tolerance to more than one abiotic stress at a time (a phenomenon known as multiple abiotic stress tolerance, or MAST). Several reports are already available in wheat where a single gene/QTL is known to provide tolerance to two or more abiotic stress^14,15,16,17^.

Considering this, it would be fascinating if these already identified QTLs could be presented in an integrated manner with a comprehensive analysis to evaluate them. To do so, a systematic, statistical strategy that takes into consideration the parameters used in previous experiments is needed to generate reliable, robust, and stable target QTLs. However, the pyramiding of multiple QTLs for distinct stresses can be undertaken by breeders in an effort to breed for MAST. However, because this strategy starts with integrating two QTLs for two different stresses into one line and then adding another QTL to get the desired traits once the generation is established, it can be a lengthy procedure. Furthermore, the introgressed QTLs may interact epistatically, affecting the outcome of the introgression process. The attempt to introgress several QTLs can be avoided if there are instances where QTLs for different stresses coincide. To see if there is any co-localization of QTLs, all of the QTLs reported for the stresses should be presented in an integrated manner and mapped out. Meta-QTL analysis can efficiently integrate information from a multitude of QTL mapping studies, allowing for a high level of statistical power over a large amount of data^18,19,20^.

The construction of a consensus map and subsequent projection of QTL allows for the identification of regions where the QTL for a given trait is densely populated. In addition, new chromosomal positions are obtained that agree with all the maps reported in earlier studies. The decrease of confidence intervals (CIs) in MQTLs, which leads to increased genetic resolution for marker-assisted breeding (MAB) and the identification of high-confidence candidate genes (CGs), is another advantage of this approach. For instance, meta-analysis of QTLs associated with several traits has already been conducted in wheat, for instance, disease resistance^21,22.23^, nitrogen use efficiency and root related traits^24,25^, yield and associated traits^26,27^, and quality traits^28,29^. In addition, meta-analysis of QTLs associated with some of the individual abiotic stresses has also been conducted previously, for instance, (i) salinity stress^30^, (ii) drought stress^31,32^, heat stress^31,33^ and pre-harvest sprouting^72^. However, no meta-analysis for QTLs associated with WS and AS has been reported in the literature. Further, a comprehensive study covering all the major abiotic stresses has never been conducted on wheat and other cereals (except barley;^34,35^).

In addition, with the advancement of next-generation sequencing technology, high-throughput genotyping based on SNP arrays, and genome-wide association studies (GWAS), the identification of significant effects of genomic loci on complex quantitative traits is becoming increasingly popular^36^. Several studies have shown that a combination of QTL meta-analysis and GWAS may be used to investigate key genomic regions and the genetic basis of major quantitative traits^37,26^. Overall, the goal of this meta-analysis was to integrate QTLs conferring tolerance against different abiotic stresses, including DS, HS, SS, WS, PHS, and AS in wheat, in order to identify consensus genomic regions and their verification using GWAS that can be used in MQTL-assisted breeding and to compile comprehensive information for increasing tolerance against multiple abiotic stresses. The genes available in MQTL regions were identified and functionally characterized. In addition, multiple abiotic stress-responsive genes showing differential expressions in the wheat tissues were discovered using RNA-seq and microarray datasets. In addition, ortho-MQTL analysis was also performed using the information on genomic synteny and collinearity among the cereals to assess the transferability of the generated information to other crops. The findings will be used in cereal breeding programmes to improve selection for yield potential and stability under multiple abiotic stresses.

## Results

### Salient features of QTL studies, collected QTLs and consensus map

A total of 3,102 QTLs were available from 116 interval mapping studies, comprising 66 studies for individual DS and HS or combined DS and HS, 20 studies for SS, 2 studies for WS, 2 studies for AS, and 26 studies for PHS (Table S2). In these studies, 85 bi-parental populations [including doubled haploid (DH), recombinant inbred lines (RILs), F2, and backcross populations] were used, with some populations being tested for different stresses multiple times in independent studies. The population size utilised for mapping varied from 34^97^ to 1884^96^. The average number of QTLs per abiotic stress was around 517, ranging from 86 (WS) to 917 (SS) (Fig. 1a). The collected QTLs for various stresses were distributed unevenly among the 21 wheat chromosomes, ranging from 73 on 4D to 266 on 3B, with an average of around 147 QTLs per chromosome (Fig. 1b). 37.49% (1163/3102) of the total collected QTLs were found in sub-genome A, 39.23% (1217/3102) in sub-genome B, and 23.27% (722/3102) in sub-genome D. About 93% of QTLs (as many as 2,888 QTLs) had all of the information needed for projection and meta-analysis. PVE values for different QTLs also differed significantly, with 68.89% (2,137/3,102) of total QTLs and 67.83% (1,959/2,888) of QTLs used for projection having PVE values of less than 10% (Fig. 1c). Likewise, 58.89% (1,827/3,102) of total QTLs and 61.21% (1,768/2,888) of QTLs used for projection had a CI greater than 10 cM (Fig. 1d). The consensus map had 100,614 markers (mainly SSR, RFLP, and SNPs) distributed over 6,647 cM (ranging from 91.11 cM for 4B to 446.7 cM for 7A, with an average of 316.52 cM) (Table S3). Overall, there were 16.13 markers per cM, with 4.28 markers per cM on chromosome 4D and 42.2 markers per cM on chromosome 1B. Sub-genome A had 34,680 markers over 2,576.31 cM, sub-genome B had 46,359 markers over 1,890.85 cM (highest marker density with 24.52 markers/cM), and sub-genome D had 19,575 markers over 2,179.83 cM (lowest marker density with 8.98 markers/cM).

**Fig. 1.**
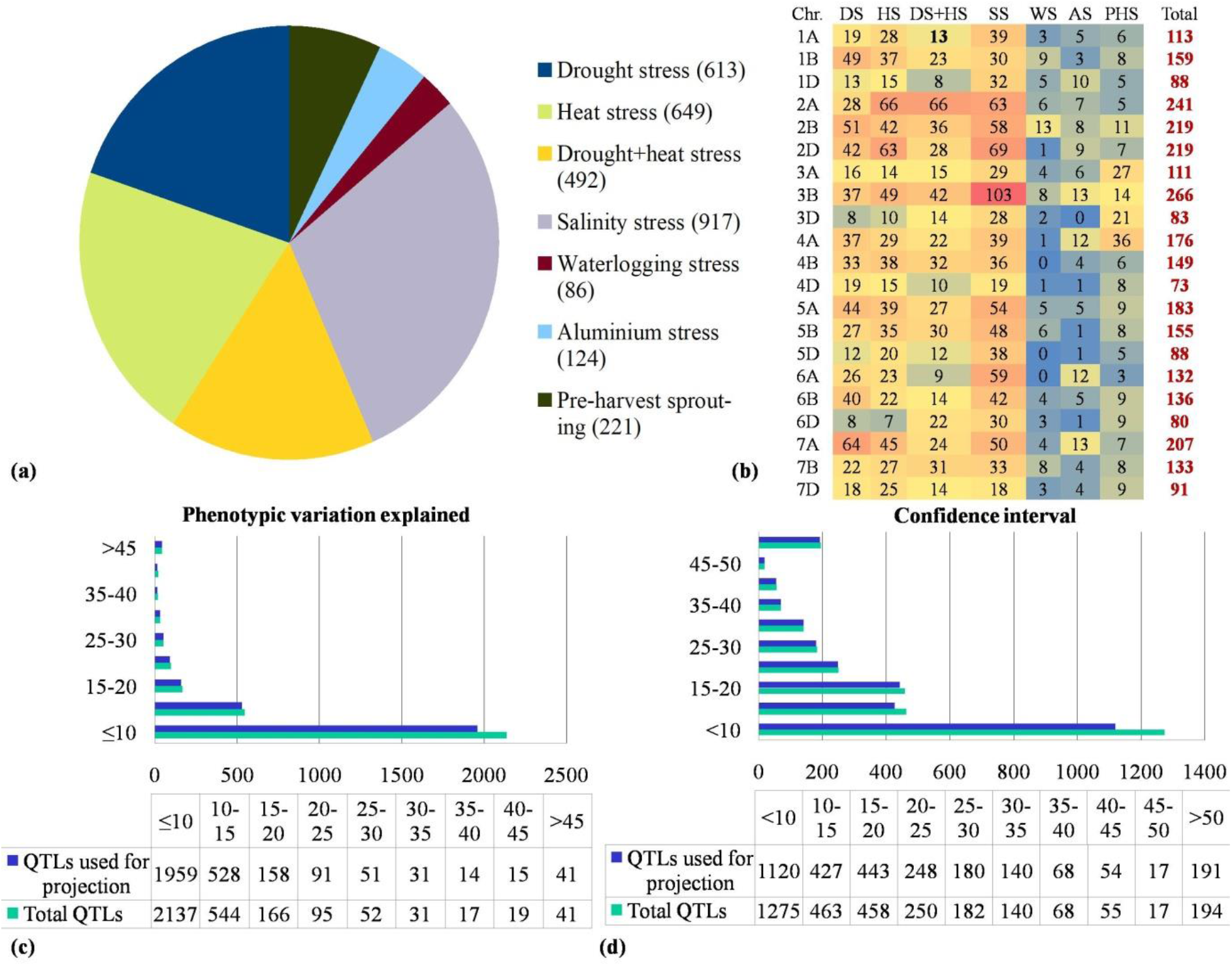
Salient characteristics of QTLs used for meta-analysis: (a) distribution of QTLs associated with different abiotic stresses, (b) chromosome-wise distribution of QTLs, (c) PVE values, and (d) confidence intervals (CI) of total collected QTLs and those used for projection

### MQTLs associated with multiple abiotic stress tolerance

Only 2,888 QTLs (out of a total of 3,102 QTLs) were utilized for projection onto the consensus map, the remaining 214 QTLs could not be used for projection because of the lack of necessary information (required for projection) from the source studies. For either of the following reasons, only 2,229 QTLs out of 2,888 could be projected onto the consensus map, leaving 659 QTLs un-projected: (i) a lack of associated markers in the consensus map, (ii) a large CI. Of 2,229 projected QTLs, 2,139 QTLs were clustered into 147 QTL hotspots, leaving 13 singletons and 47 unassigned to any hotspot. Out of 147 QTL hotspots, 13 hotspots included initial QTLs only from a single study, and therefore could not be considered as true MQTLs. The 134 true MQTLs were distributed on all 21 chromosomes, with a minimum of only one MQTL on chromosome 3A and a maximum of 10 MQTLs on chromosome 7A (Table S4; Fig. 2a). The number of QTLs included in an individual MQTL ranged from only 2 QTLs in seven MQTLs (MQTL3D.5, MQTL4D.6, MQTL5B.1, MQTL6A.2, MQTL6A.8, MQTL6B.1, MQTL6B.7) to a maximum of 107 QTLs in MQTL3B.4. In addition, 45 MQTLs were based on fifteen or more QTLs from different studies (Fig. 2b). Furthermore, the number of MQTLs did not match to the number of initial QTLs on individual chromosomes. For instance, the proportion of MQTLs on chromosomes 2A and 3A was low in comparison to the number of QTLs involved.

**Fig. 2.**
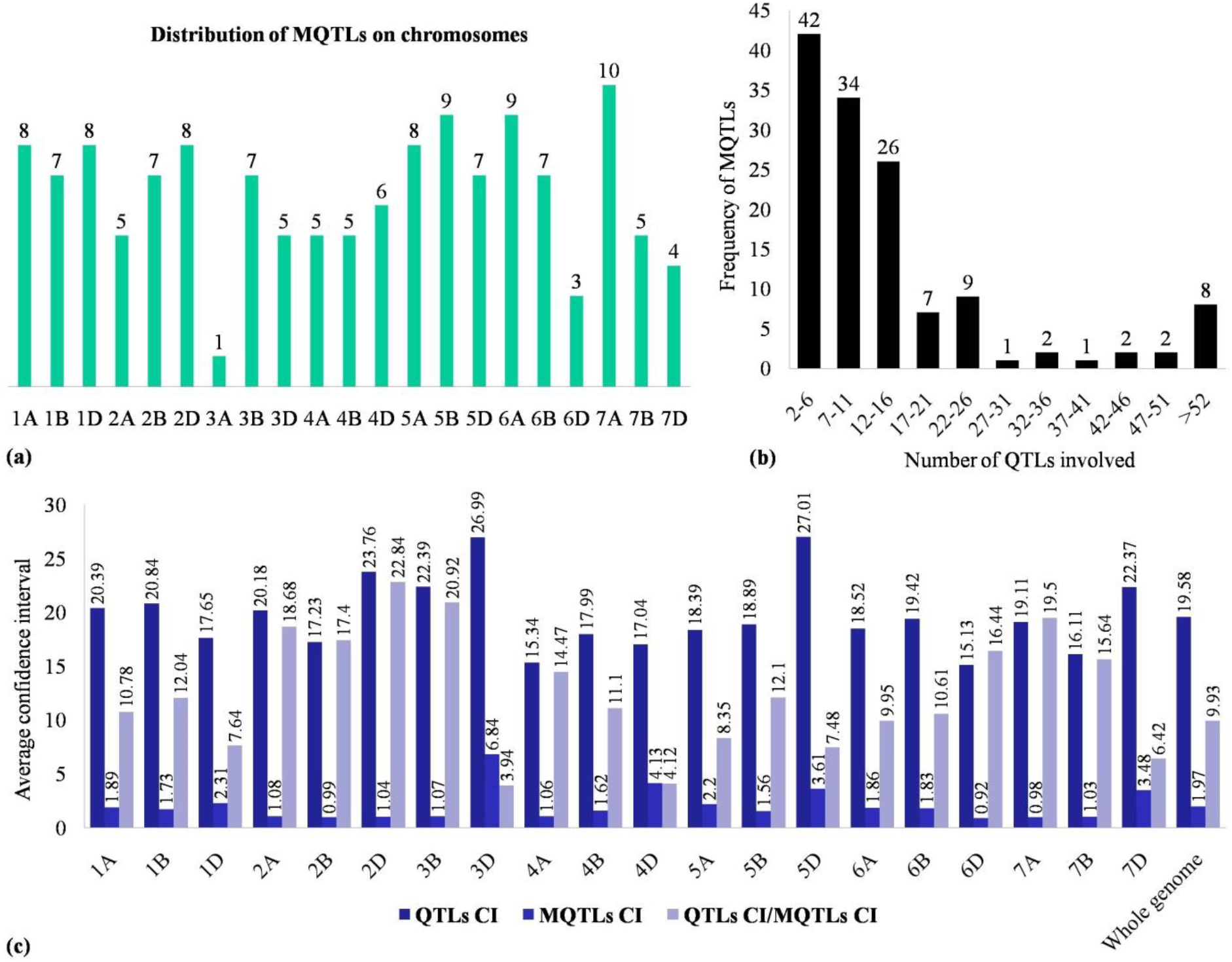
Salient features of MQTLs detected in the current study: (a) chromosome-wise distribution of MQTLs, (b) frequency of QTLs included in different MQTLs, and (c) comparison of CIs of initial QTLs and MQTLs

Individual MQTLs explained 2.9 to 36.06% of phenotypic variation, whereas LOD scores ranged from 2.81 to 33.32 %. The MQTLs had an average CI of 9.93 times smaller than the initial QTLs (the average CI of initial QTLs was 19.58 cM), and there were substantial differences in CI reduction among different chromosomes. On chromosomes 2D and 3B, the average CI of MQTL was reduced by 22.84 and 20.92 times, respectively, followed by 19.5 and 18.68 times on chromosomes 7A and 2A, respectively (Fig. 2c). Interestingly, only a single MQTL was identified on chromosome 3A, which showed a CI of only 0.01 cM, whereas, the initial QTLs located on chromosome 3A had an average CI of 16.35 cM. Further, all MQTLs (except MQTL5B.9 and MQTL5D.7) were physically positioned on the wheat reference genome. The physical CI ranged from 4,553 bp (MQTL6B.6) to 653.8 Mb (MQTL6B.1) with a mean of 47.09 Mb. In total, 72% (97/134) of MQTLs occupied a physical length of less than 20 Mb in the wheat genome. There were five MQTLs (MQTL2B.3, MQTL3A.1, MQTL3B.4, MQTL5A.2, and MQTL7B.3) that provided tolerance against all the six stresses; 23 MQTLs each conferred tolerance to five different stresses (Table 1); the remaining MQTLs provided tolerance to 1 to 4 abiotic stresses.

### MQTL validation and the availability of known genes within MQTL regions

There were considerable overlaps between the MQTLs discovered through meta-analysis and the significant markers (MTAs) identified through the GWAS approach that are associated with various abiotic stresses in the wheat genome. Remarkably, 817 MTAs identified in wheat GWAS for various abiotic stresses overlapped with 76 MQTLs out of the 132 physically anchored MQTLs (Table S5, Fig. 3). For instance, there were significant differences in the number of MTAs overlapping with an individual MQTL. For instance, the MQTL2B.3 overlapped with 132 MTAs, followed by the MQTL6B.1 which overlapped with 122 MTAs, while, several MQTLs each overlapped with only one MTA. Some MQTLs (e.g., MQTL1B.3, MQTL2B.3, MQTL2D.1, MQTL4B.2) overlapped with more than 24 MTAs and involved 20 or more initial QTLs. Overall, GWAS was used to validate 57% of the total MQTLs.

**Fig. 3.**
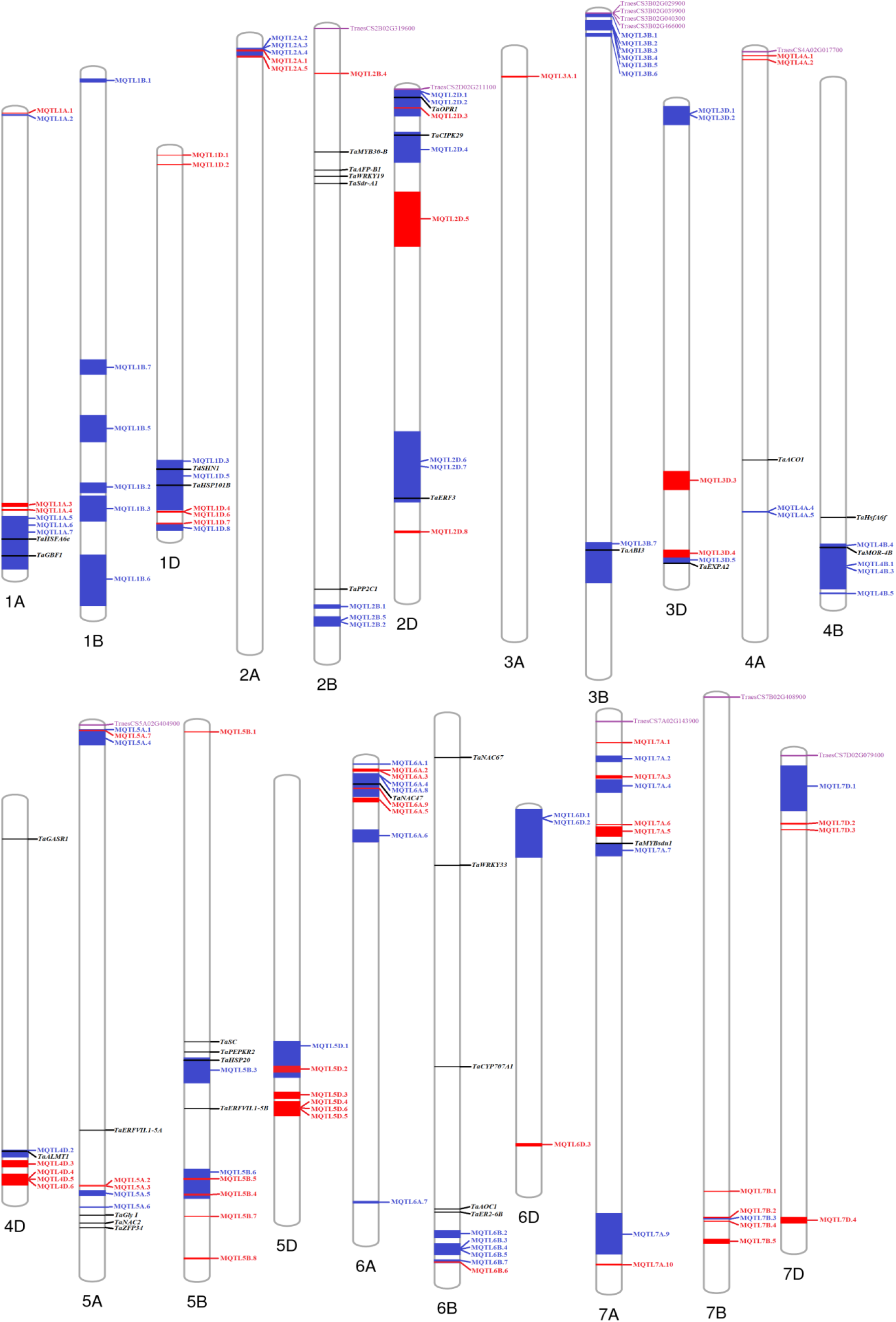
Distribution of MQTLs (GWAS-validated MQTLs are shown with blue), differentially expressed candidate genes (showing differential expression in at least three datasets), and known genes co-localizing with MQTLs (some of the MQTLs with large physical regions are not shown).

The MQTLs identified in the current study also co-localized with 34 known genes conferring tolerance to various abiotic stresses [including 5 genes for DS (*TaERF3, TaMOR-4B, TaZFP34, TaER2-6B*, and *TaWRKY33*), 6 for HS (*TaHSFA6e, TaHSP101B, TaHsfA6f, TaGASR1, TaPEPKR2*, and *TaHSP20*), and 16 for SS (*TaGBF1, TdSHN1, TaWRKY19, TaPP2C1, TaMYB30-B, TaOPR1, TaCIPK29, TaEXPA2, TaACO1, TaGly I, TaNAC2, TaSC, TaNAC47, TaAOC1, TaNAC67*, and *TaMYBsdu1*), 2 for WS (*TaERFVII*.*1-5A* and *TaERFVII*.*1-5B*), 1 for AS (*TaALMT1*), and 4 for PHS (*TaSdr-A1, TaAFP-B1, TaABI3*, and *TaCYP707A1*). For instance, MQTL2B.6 overlapped *TaSdr-A1, TaAFP-B1, TaWRKY19*, and *TaMYB30-B* genes, MQTL5A.8 overlapped *TaZFP34, TaGly I, TaNAC2*, and *TaERFVII*.*1-5A* genes, MQTL5B.2 co-localized with *TaPEPKR2, TaSC*, and *TaERFVII*.*1-5B* genes, and MQTL6B.1 overlapped *TaER2-6B, TaWRKY33, TaCYP707A1, TaAOC1*, and *TaNAC67* genes (Table S6, Fig. 3). These genes encode different types of proteins involved in different pathways, for instance, transcription factors (e.g., AS2/LOB, ERF, MYB, NAC, and WRKY), protein kinases (e.g., CBL-interacting protein kinase, LRR receptor-like kinase protein, phosphoenolpyruvate carboxylase kinase), transporters (e.g., aluminium-activated malate transporter) and other proteins including ABA 8′-hydroxylase, ABI5 binding protein, allene oxide cyclase, aminocyclopropane-1-carboxylate oxidase, GA-stimulated transcript family protein, glyoxalase, nicotianamine synthase, etc.

### Candidate genes available from MAST-MQTLs

As many as 539 gene models were detected in the genomic regions of 28 selected MQTLs, each based on at least nine QTLs (providing tolerance to at least five different abiotic stresses) and with an average genetic and physical CI of 0.65 cM and 59.66 Mb, respectively (Table S7). These gene models included 66 gene models with no function descriptions. At the two extremes, a maximum of 45 gene models were available from MQTL1D.4, whereas, a minimum of only 4 gene models were available from MQTL2B.3. Gene models with similar protein products were identified repeatedly in different MQTL regions, for instance: 27 gene models for proteins with protein kinase domains, 18 for TRAF-like proteins, 14 for proteins with F-box domains, 14 for zinc finger proteins (including C2H2-type, Dof-type, GRF-type, RING-type), 9 for nucleotide-diphospho-sugar transferases, 7 gene models each for cytochrome P450 proteins, for proteins belonging to the glycoside hydrolase family, and for UDP-glucuronosyl/UDP-glucosyltransferases, 5 gene models for ABC transporter-like proteins, etc.

### In silico expression analysis of genes and common cis-regulatory elements in the promoters of multiple abiotic stress responsive genes

In silico expression analysis was performed on all 539 gene models available from 28 selected MQTLs. The first transcriptomic dataset (which included data on genes expressed under DS, HS, and combined DS and HS conditions) identified 70 differentially expressed genes (DEGs), including 25 up-regulated genes, 7 down-regulated genes, and 38 genes that were up-regulated in some conditions but down-regulated in others. MQTL7D.1 had the most DEGs (7), but none were available for MQTL2A.5, MQTL3B.6, MQTL7A.5, MQTL7A.6, MQTL7B.3, and MQTL7D.3. The second dataset (which included gene expression data under PEG-simulated drought conditions) identified 81 DEGs, with 7 genes up-regulated, 44 genes down-regulated and 30 genes up-regulated at the one time point but down-regulated at another. From this dataset, MQTL4A.5 had a maximum of 11 DEGs, whereas, no DEG was available from the following five MQTLs: MQTL2A.5, MQTL3A.1, MQTL7A.5, MQTL7A.6, and MQTL7D.3. The third dataset (including data on differential expression of genes in a DH population grown under water stress conditions) uncovered a total of 108 DEGs with 52 up-regulated and 56 down-regulated genes. The fourth dataset (which contained data on genes expressed during grain development with and without the dormancy QTL) contained a maximum of 10 DEGs; no DEGs were obtained from the MQTL2A.5 and MQTL7B.3. Similarly, MQTL3B.4 had a maximum of 4 DEGs, whereas, several MQTLs gave no DEGs.

Overall, 191 DEGs were identified from 27 MQTLs (no DEG was available from MQTL2A.5). Furthermore, 27 genes were common between the first and second transcriptomic datasets, and 36 genes were common between the third and fourth transcriptomic datasets. There were 11 genes (*TraesCS2B02G319600, TraesCS2D02G211100, TraesCS3B02G029900, TraesCS3B02G039900, TraesCS3B02G040300, TraesCS3B02G466000, TraesCS4A02G017700, TraesCS5A02G404900, TraesCS7A02G143900, TraesCS7B02G408900, TraesCS7D02G079400*) which were common among first, second and third datasets; whereas, two genes (*TraesCS2D02G211100, TraesCS3B02G039900*) showed differential expression across all the four transcriptomic datasets investigated (Table 2). A representative heat map of these 11 most promising DEGs is presented in Fig. 4.

**Fig. 4.**
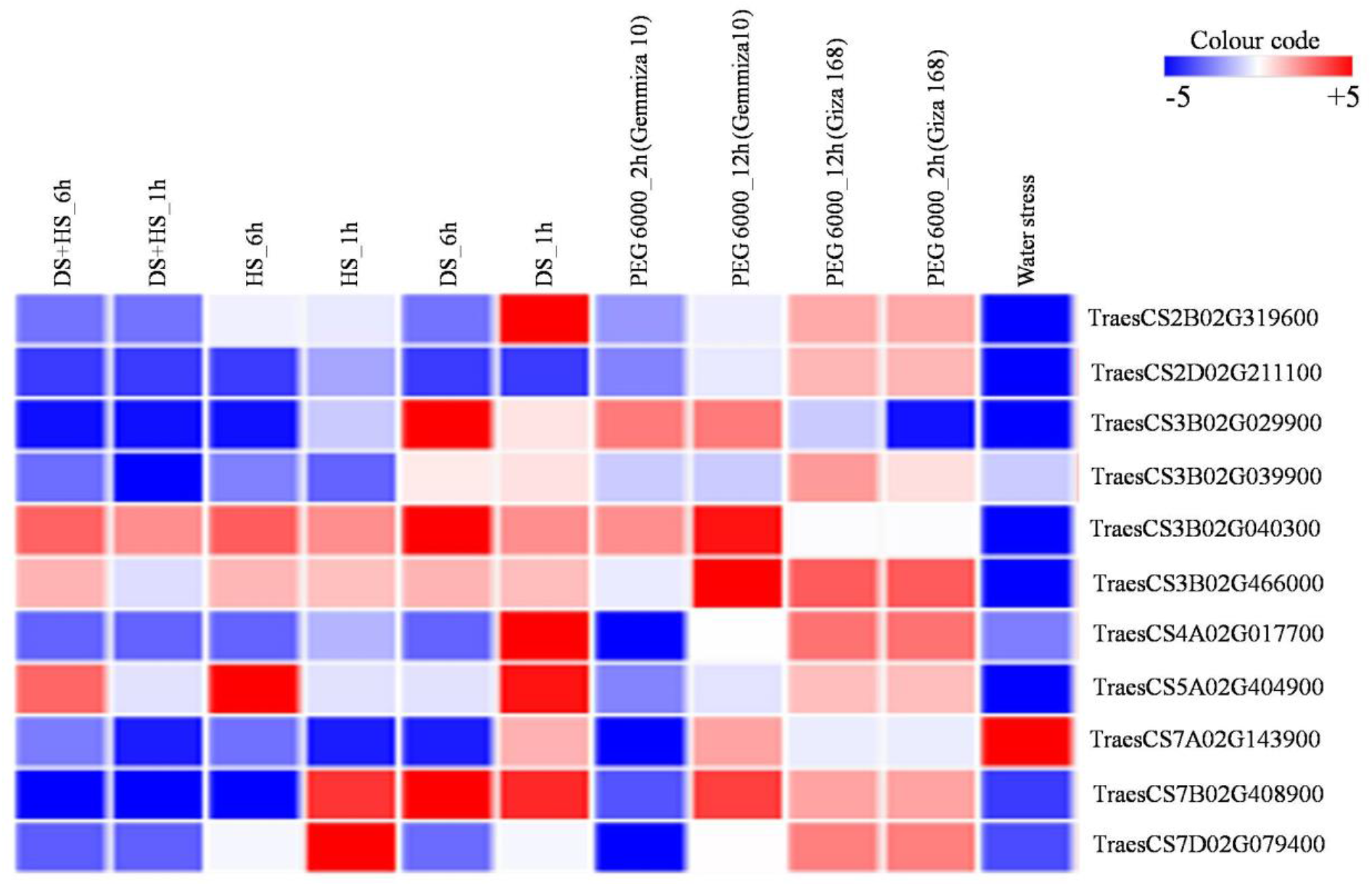
Differential expression of 11 most promising candidate genes under different conditions.

Gene control is mediated by cis-regulatory elements (CREs) in promoter regions. It is crucial to study the CREs found in the putative promoter regions of the DEGs reported in this study. CREs were studied in 1.5 kb 5′ upstream promoter regions of each of the 11 multiple abiotic stress-responsive genes. A total of 374 CREs were discovered, including 20 unnamed/uncharacterized CREs (Table S8). The CREs were divided into three categories: plant growth and development, stress responsiveness, and phytohormone responsiveness, in addition to the conventional cis-elements found in promoters of DEGs (Fig. 5). Shoot and root meristem expression (CAT-box), endosperm expression (GCN4-motif and AAGAA-motif), flowering (CCAAT-box, AT-rich element, and Circadian), and zein metabolism (O2 site) were all included in the plant growth and development category. Drought-inducibility (MBS), anaerobic induction (ARE), stress (TC-rich repeats, As-1 element, STRE), low temperature (LTR), and wounding responsiveness (WRE3 and WUN-motif) were all included in the stress responsiveness group. Many cis-elements, including salicylic acid (TCA-element), ethylene (ERE), Me-JA (TGACG-motif and MYC), auxin (TGA-element), and abscisic acid (ABRE), were shown to be related to phytohormone responsiveness. ABRE, MYC, and ERE were the most common cis-elements, and they were all related to hormone responsiveness. These findings show that complex regulatory networks play a role in the transcriptional regulation of genes that play a role in MAST.

**Fig. 5.**
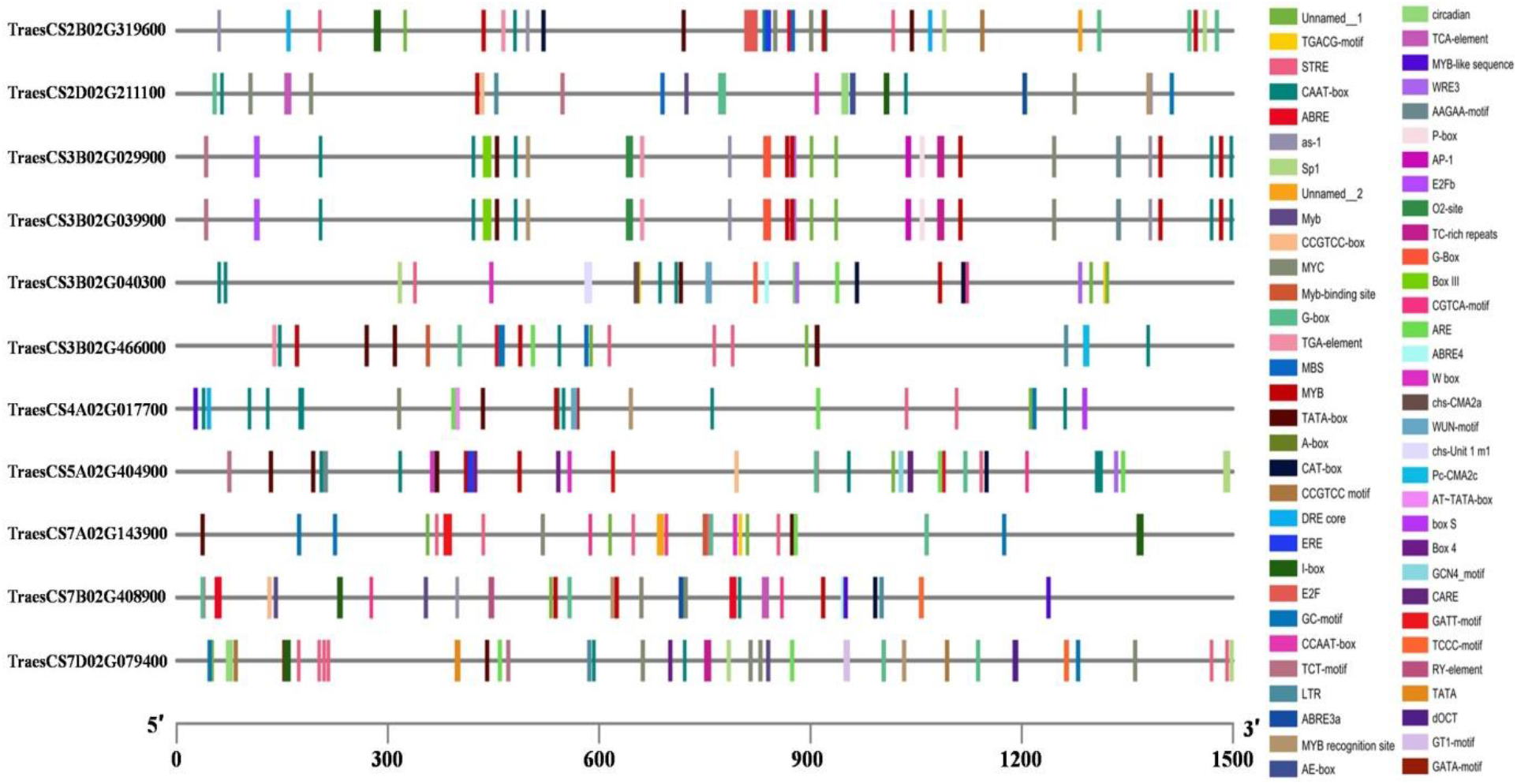
Information on cis-regulatory elements in promoter regions of 11 multiple abiotic stress tolerance genes

### Conserved genomic regions control tolerance to multiple abiotic stresses in different cereals

The syntenic relationship was also observed for wheat genes available from MQTL regions with the maize and rice genomes. The 5,161 and 5,893 wheat genes available from MQTLs were used to detect the 5,842 maize orthologues, and 5,277 rice orthologues, respectively (Fig. 6a, b). This synteny analysis revealed several genomic regions in both maize and rice genomes where several MQTLs have already been identified in previous studies for different abiotic stresses; such genomic regions conserved between ‘wheat and maize’ and ‘wheat and rice’ may be termed as ‘ortho-MQTLs’. Out of 5,842 maize genes, 2,565 genes were available in 101 maize MQTL regions previously identified to be associated with drought stress tolerance^38,39,40^. Similarly, out of 5,277 rice genes, 2,704 genes belonged to 140 rice MQTLs earlier reported to be associated with heat, drought, and salinity stress tolerance^41,42,43,44,45^. The number of conserved genes at corresponding orthologous positions of MQTLs (between ‘wheat and maize’ and ‘wheat and rice’) ranged from just a few to hundreds of genes. Detailed information related to maize and rice orthologues of wheat genes and ortho-MQTLs is presented in Table S9. Some earlier studies^46,35^ did not provide physical positions of MQTLs; thus, these studies could not be used to mine ortho-MQTLs.

**Fig. 6.**
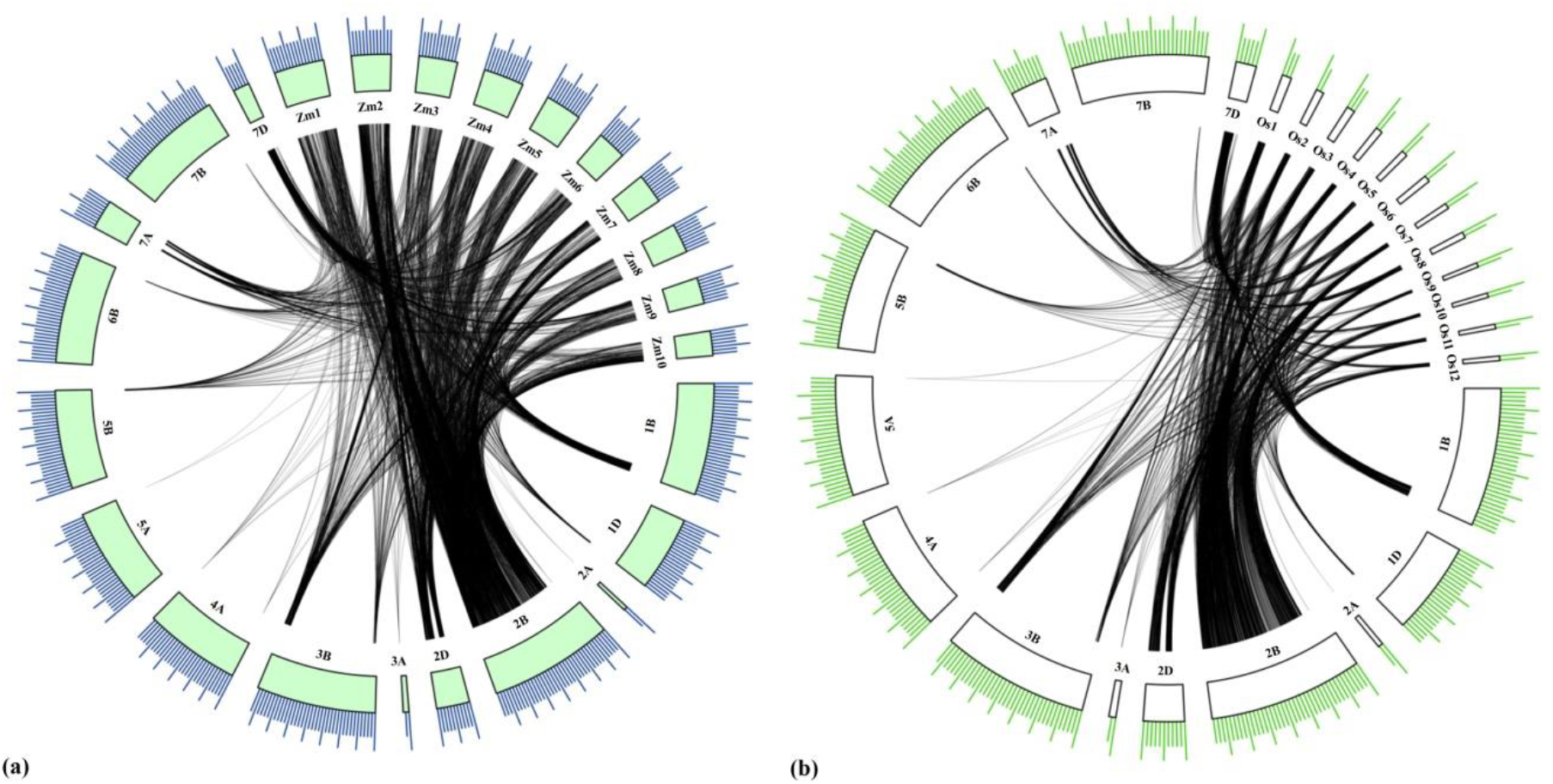
Syntenic relationship of genes available from the wheat MQTL regions with the (a) maize and (b) rice genomes. The physical sizes of the chromosomes are indicated by the ruler drawn above on them, with larger and smaller tick marks at every 100 and 20 Mbp, respectively.

## Discussion

DS, HS, SS, WS, PHS, and AS are all abiotic stresses that are detrimental to wheat, posing significant intimidation to the environment and agriculture and resulting in considerable crop yield loss. These abiotic stresses may be avoided by growing wheat varieties conferring tolerance to individual or multiple abiotic stresses. Thus, new wheat cultivars must contain novel loci/genes to provide tolerance against these stresses that influence plant development and yield. A number of studies on QTL mapping for different abiotic stresses in wheat have been undertaken over the last two decades (Table S1). Most of the QTLs found in these previously reported studies had a long CI and a relatively low PVE, which made them ineffective for use in MAB to develop wheat varieties providing tolerance against different abiotic stresses. Furthermore, validation is also required whenever these QTLs are planned to be used in a wheat breeding programme. In comparison to the QTLs that have been previously identified, MQTLs are observed to be more stable and consistent, with a narrow CI and comparatively higher confidence, making them useful for crop improvement programmes as well as for basic research programmes, including the cloning and further characterization of favorable QTLs/genes^26^. Furthermore, several QTLs have been detected and characterized, each conferring tolerance to multiple abiotic stresses in different crops, including wheat^47,48,49,10,50,51,52^. Furthermore, numerous studies reported the availability of individual genes conferring tolerance to multiple abiotic stresses in wheat, which are regarded as extremely valuable assets for future breeding programmes aimed at developing climate-smart wheat varieties^15,53,14^. Based on these studies, the MAST hypothesis was figured out, which says that one locus is thought to make a plant more resistant to more than one abiotic stress.

Several meta-QTL analyses for various traits in cereal crops such as maize^39^, rice^41,46,20^, and barley have already been conducted^35^. In wheat, meta-QTL studies have been undertaken for four different abiotic stresses, including PHS^54^, SS^30^, DS^32^, and HS^33^, but no meta-analysis has been conducted for the traits contributing to WS and AS stresses. Furthermore, to our knowledge, no MAST meta-analysis has ever been performed on wheat. In addition, the information from such studies quickly becomes outdated due to the identification and characterization of new QTLs in recently conducted studies. This necessitates a comprehensive meta-QTL analysis at a regular interval for proper utilization in the breeding pipeline. Meta-QTL studies for abiotic stresses in wheat have been done before. In this study, we looked at the QTLs for six different abiotic stresses that have been reported up until 2021).

This study’s consensus map is much better and more comprehensive than previous consensus maps used for MQTL for some of these abiotic stress tolerance traits^32,33,30^. The consensus map contained 100,614 markers (mostly SSR and SNPs) scattered throughout 6,647 cM. Based on the QTL information and the development of the consensus map in this study, it is observed that the sub-genome B carries the maximum number of markers and contains the highest density of QTLs, while the sub-genome D has the minimum number of markers and contains the lowest density of QTLs. The characteristics of marker distribution and density of QTLs on different wheat chromosomes analysed in the current meta-study are in agreement with other meta-studies on individual abiotic stress tolerance earlier conducted in wheat^31,32,33^.

In this study, we predicted 134 MQTLs using previously identified QTLs. The majority of MQTLs (87.31%; 117/134) were each associated with tolerance against at least two of the six stresses under study. Furthermore, 23 MQTLs each were involved in tolerance for at least five different stresses, while five MQTLs, each were associated with all six abiotic stresses under study. As discussed, several MQTLs contained QTLs for different abiotic stress tolerance traits, suggesting a strong association between different abiotic stress tolerance traits. Furthermore, QTLs for the same abiotic stress tolerance from different mapping populations were projected onto the same chromosomal region, confirming that those regions exist. The meta-analysis conducted in this study refined the CIs and compared the genomic positions of different QTLs detected in individual studies associated with different stresses. The average CI of MQTLs in this study was reduced by 9.93 times compared to the CI of initial QTLs. A circular diagram representing the salient characteristics of QTLs and MQTLs is provided in Fig. 7.

**Fig. 7.**
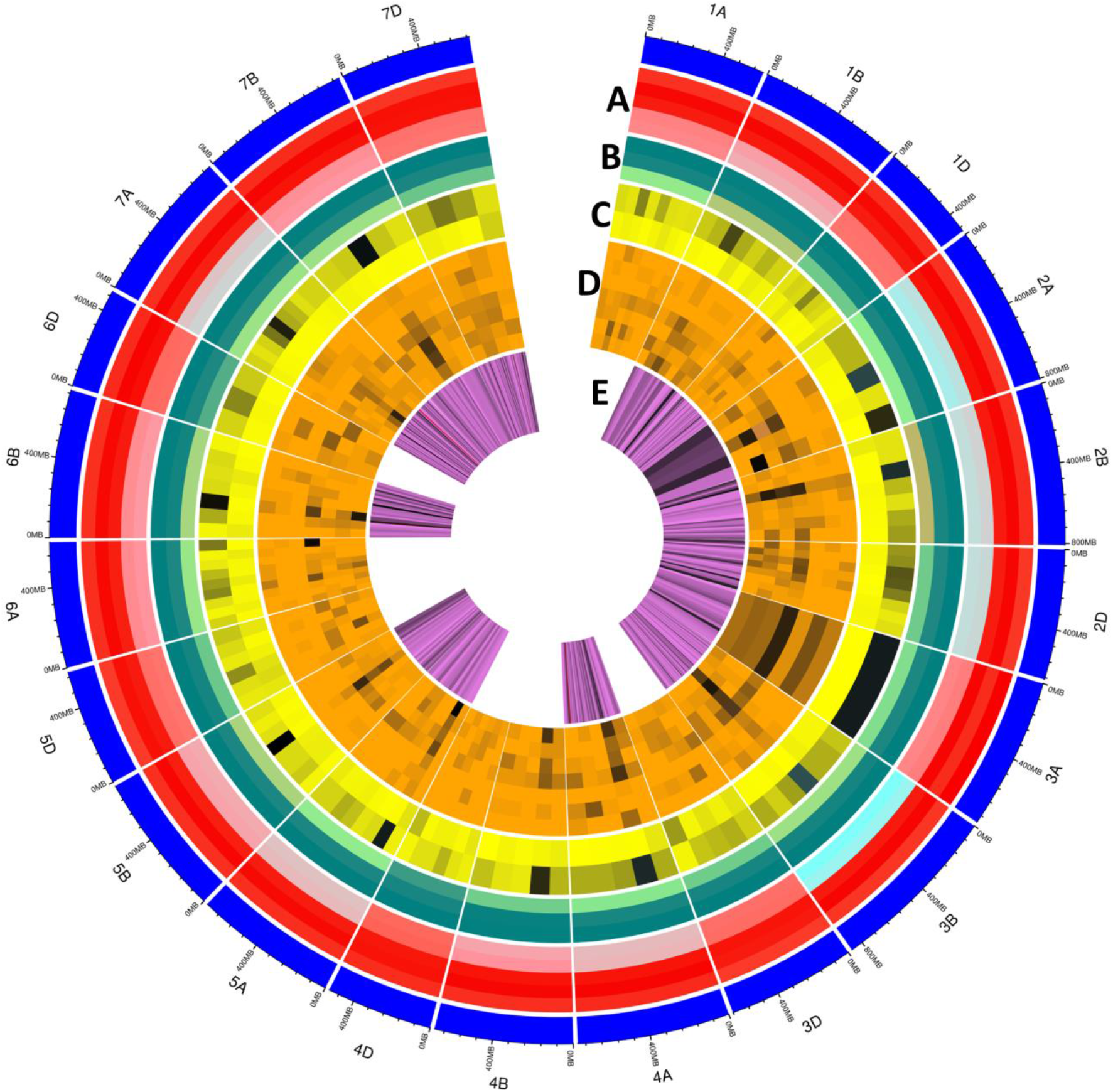
Circular diagram depicting the salient characteristics of QTLs and MQTLs. Circle A- chromosome wise distribution of number of QTLs, QTLs projected on consensus map, average confidence interval (CI) of MQTLs and average CI of initial QTLs. Circle B- Number of markers, chromosome lengths and density of markers on different chromosomes. Circle C- The number of QTLs involved in individual MQTL. Circle D- The number of QTLs for each stress in individual MQTLs (i.e., DS, HS, DS+HS, SS, WS, PHS, and AS) and number of traits associated with individual MQTLs. Circle E- Densities of the gene in different MQTL regions. The physical maps of wheat chromosomes are represented by the outer circle.

Interestingly, more than 57% (76/134) of MQTLs were validated with GWAS-MTAs associated with different abiotic stress tolerance traits identified in wheat. The MQTL2B.3 overlapped with the maximum number of MTAs (132) obtained from multiple GWA studies, while some MQTLs overlapped with only one MTA. Some recent studies also provided similar results, such as 54.6%, 63%, and 61.3% verification of MQTLs with GWAS, respectively^23,26,27^. Furthermore, the MQTLs validated using GWAS data are important genomic regions that could be utilized effectively in future wheat breeding programmes and cloning of MAST genes available from MQTLs.

A comparison of MQTLs with known genes for distinct abiotic stresses in wheat could aid investigation into the molecular mechanisms that confer tolerance to various abiotic stresses. As a result, the relationships between MQTLs and known genes were also investigated in the present study. Twenty-one MQTLs were found to be co-localized with 34 genes in wheat that are involved in providing tolerance to different abiotic stresses. For instance, MQTL2B.6 overlapped with 2 PHS tolerance genes (*TaSdr-A1*^35^ and *TaAFP-B1*^55^), and 2 SS genes (*TaWRKY19*^56^ and *TaMYB30-B*^57^). MQTL5A.8 overlapped with one DS gene (*TaZFP34*^58^), two SS genes (*TaGly I*^59^ and *TaNAC2*^60^), and one WS gene (*TaERFVII*.*1-5A*^61^). MQTL5B.2 co-localized with one HS gene (*TaPEPKR2*^14^), one each for SS (TaSC^62^ and WS (*TaERFVII*.*1-5B*^61^). Similarly, MQTL6B.1 overlapped with two DS genes (*TaER2-6B*^63^ and *TaWRKY33*^64^), one PHS gene (*TaCYP707A1*^65^), and two SS genes (*TaAOC1*^66^ and *TaNAC67*^17^). According to these findings, the twenty-one MQTLs, which overlapped with 34 known genes for different abiotic stresses, appear to be promising candidates for improving tolerance to multiple abiotic stresses and could be valuable assets in breeding for climate resilience. However, more work is required to find functional variants of these genes in the MQTL regions that are being studied.

Furthermore, there are several genes known in wheat that confer tolerance to multiple abiotic stresses^35,67,16,60,68^. Unfortunately, no MAST gene or QTL providing tolerance against all the six abiotic stresses under this study has been identified. In the present study, mining of genes in genomic regions of the 28 selected MAST-MQTLs allowed the identification of 539 non-redundant gene models. *In silico* expression analysis unravelled the 11 promising CGs showing differential expressions in at least three transcriptomic datasets. These genes included 7 genes encoding for characterized proteins (viz., late embryogenesis abundant protein, sugar transporter-like, O-acyltransferase, zinc finger protein, transmembrane protein, fatty acid hydroxylase, and protein kinase), and the remaining 4 genes encode for uncharacterized proteins (viz., protein of unknown function DUF1666, protein of unknown function DUF247, uncharacterized conserved protein UCP031279, and un-characterized protein).

The relevance of some of these genes encoding for known proteins can be summarized as follows: (i) Late embryogenesis-abundant (LEA) proteins are a large and diverse family of proteins that are hypothesized to play a role in normal plant growth and development as well as protecting cells against a variety of abiotic stresses. Chen and colleagues^69^ identified and characterized several LEA family genes conferring tolerance to multiple environmental stresses^69^. (ii) In a recent study, Xuan and colleagues^70^ reported that a higher expression level of sugar transporter genes regulates plant responses to different stresses, including drought, salt, and low-temperature stress. (iii) Transgenic plants with increased expression of the *WSD1* gene, which encodes for *wax synthase/acyl-CoA:diacylglycerol acyltransferase* showed improved tolerance to abscisic acid, mannitol, drought, and salinity^71^. These findings showed that manipulating cuticular waxes could be beneficial for increasing plant productivity in a rapidly changing climate. (iv) Overexpression of *ZFP182*, a TFIIIA-type zinc finger protein significantly improved tolerance against salt, cold, and drought stresses in transgenic rice^72^. Further, zinc finger proteins have been termed the master regulators of abiotic stress responses in plants^100^. However, these genes are assumed to have a pleiotropic function, but there is still no evidence of how these genes may confer tolerance to multiple abiotic stresses; further studies are required to characterize their functionality in response to multiple abiotic stresses. Moreover, the DEGs with no function descriptions available may be characterized for their possible roles in MAST in wheat in further studies.

Since there is a cross-talk between different abiotic stresses, identification of common and/or overlapping CREs is crucial for developing wheat cultivars that show tolerance towards them. CREs are DNA sequences found in gene promoters that transcription factors bind to, and they further drive gene expression by either enhancing or silencing transcription regulations in response to unfavorable environmental conditions^73^. In the present study, all 11 multiple abiotic stress-responsive DEGs were identified to be equipped with three different functional categories of cis-elements, such as plant growth and development, stress responsiveness, and phytohormone responsiveness. In the case of stress responsiveness cis-elements, for instance, the MYB binding site (MBS) element present in the Zm*SOPro* promoter region triggers drought stress tolerance in transgenic *Arabidopsis*^74^. Stress-responsive gene regulation in *Arabidopsis* and soybean essentially depends on the TGGTTT (ARE), activation sequence-1 (as-1) and TC-rich repeat elements^75,76,77^. In addition, WUN-motif is a dehydration-related element detected in the regulatory region of the *StDREB* gene in tomato^78^.

The current study additionally investigated chromosomal regions with MQTLs that are conserved across several cereals. These ortho-MQTL regions were identified by integrating the two different approaches utilized by Saini et al.^26^ and Sandhu et al.^20^, respectively. Ortho-MQTLs have recently been conducted in wheat for different traits including quality traits^28^, thermotolerance^33^, salinity stress tolerance^30^, nitrogen use efficiency^24^, grain yield and component traits^22^. The discovery of ortho-MQTLs in closely related species shows their relevance and stability, as well as the reliability of associated genes^54^. Functional validation of the orthologous genes uncovered in this study is feasible, and gene-based functional markers for cereal breeding programmes can be developed^79^.

Further, based on the criteria given in Saini et al.^26^, we selected the five most promising MQTLs for MAST breeding programmes in wheat (viz., MQTL1D.4, MQTL2D.5, MQTL3A.1, MQTL4A.2, and MQTL7A.6). These MAST-MQTLs, which are involved in providing tolerance to at least five abiotic stresses under study, had CI ranging from 0.01 to 1.16 cM and comparatively high PVE values (ranging from 12.19 to 21.66). In addition, these MQTLs included 16-79 initial QTLs derived from 11 to 35 interval mapping studies. Therefore, these are believed to be the most stable and consistent MAST-MQTLs across all genetic backgrounds and environments. These MAST-MQTLs are important breeding materials. They can be used either in MAB programmes aimed at developing wheat varieties providing tolerance to multiple abiotic stresses under study or subjected to fine mapping to identify key genes involved in tolerance against multiple abiotic stresses in wheat.

### Concluding remarks

The results of interval mapping studies on DS, HS, SS, WS, PHS, and AS tolerance were successfully integrated during the current study, resulting in the identification of 134 MQTLs. GWAS was used to confirm more than half of these MQTLs. Thirty-four known genes associated with various abiotic stresses overlapped several of these MQTLs as well. Furthermore, 28 MQTLs conferred tolerance to five or all of the six abiotic stresses studied. Putative genes associated with several abiotic stresses were discovered, and 11 genes encoding important proteins showed differential expression across different transcriptome datasets. For the development of tolerance to numerous abiotic stresses in wheat cultivars, five promising MQTLs imparting tolerance to at least five abiotic stresses were recommended for use in marker-assisted breeding. Overall, these findings provide useful information as well as robust candidate genes for further functional investigation aimed at enhancing wheat MAST.

## Material and methods

### Collection of QTL mapping studies and data on QTLs

The literature associated with QTL mapping studies of different abiotic stresses, including DS, HS, SS, WS, PHS, and AS in wheat, was collected using relevant keywords through online platforms such as PubMed (https://www.pubmed.ncbi.nlm.nih.gov/) and Google Scholar (https://www.scholar.google.com/). This investigation was supplemented by a newly developed wheat QTL database (WheatQTLdb; www.wheatqtldb.net;^98^. Each independent study was further used to obtain the following information: (i) markers flanking the QTLs, (ii) information on mapping populations (size and type), (iv) genetic peak position and confidence interval (CI), (v) LOD score, and (vi) phenotypic variation explained by individual QTLs (R2 or PVE).

Studies with incomplete datasets were excluded. In some cases, the peak position of the QTL was not available, consequently, the mid-point of both the flanking markers was considered the peak. Simultaneously, a LOD score of 3.0 and a PVE value of 10 were assigned wherever no LOD score or PVE value was provided. All of the QTLs involved in this study were given unique identities as follows: the author’s name followed by the year of publication, the abbreviated name of the trait, stress, and chromosome involved, respectively. Numerals after the chromosome numbers were used to distinguish various QTLs for the same trait and stress on the same chromosome. Under each stress condition, QTLs associated with different traits, including yield and related traits, and several morpho-physiological traits, were reported in the literature. For the purpose of this analysis, all the QTLs associated with different component traits identified under a particular stress condition were treated as QTLs associated with the abiotic stress in question.

### Construction of the consensus map and projection of QTLs

A highly dense consensus map was constructed by integrating the following high-quality linkage maps: (a) ‘Wheat, Consensus SSR, 2004’^80^ (b) ‘*Synthetic x Opata BARC* map’^81^ and SNP maps assayed through (c) ‘Illumina iSelect 90K SNP Array’^82^, (d) ‘Wheat 55K SNP array’^83^. In addition, markers flanking the individual QTLs were also integrated into this consensus map to ensure the projection of the maximum number of QTLs. The R package ‘LPMerge’ was utilized to generate the consensus map^84^.

Separate map and QTL data input files from several QTL mapping experiments were prepared. The map file included information mainly on linkage groups, markers, and their genetic positions, whereas, the QTL file contained information on QTL name, stress name, chromosome number, LOD, PVE, and genetic positions (peak position and CI) of individual QTLs. In some studies where the original CI for individual QTLs was not reported, CI (95%) was obtained for each QTL using different population-specific equations^85,86,87^. According to Chardon et al.,^99^ QTLs were projected onto the newly created consensus map using the projection tool (QRL-Proj) in Biomercator v4.2 software (2004). To select the optimum projection context, QTLProj employs a dynamic strategy. A perfect context consists of a pair of common markers that flank the QTL in the original map and a consistent distance estimate between the two maps (initial and consensus). The behavior of the procedure to get such a configuration is influenced by the minimal value of the flanking marker interval distances ratio and the minimal p-value produced by confirming the homogeneity of the flanking marker interval distances between the initial and consensus maps^19^.

### Prediction of MQTLs and their validation using GWAS

Individual MQTLs were predicted for each wheat chromosome as per instructions provided in the manual (https://www.ebi.ac.uk/eccb/2014/eccb14.loria.fr/programme/id_track/ID10-summary.pdf) and keeping abiotic stress as a meta-trait (including all six abiotic stresses: DS, HS, SS, WS, PHS, and AS). The two-steps method proposed by Veyrieras et al.^19^ was utilized for the prediction of MQTLs. The first step involves the selection of the best MQTL model based on obtaining the lowest values of the selection criteria in at least three of the following five models: (i) AIC (Akaike information content), (ii) AICc (AIC correction), (iii) AIC3 (AIC 3 candidate models), (iv) BIC (Bayesian information criterion), and (v) AWE (average weight of evidence). The second step predicts the total number of MQTLs on a chromosome along with information about peak position, CI (95%), and weightage based on the model selected. An MQTL was described as a genomic region in which two or more QTLs from distinct mapping studies were involved. The means of the LOD and PVE values of the initial QTLs involved were used to compute the LOD and PVE values of the predicted MQTLs. When more than one MQTL was found on a single chromosome, the MQTLs were named using the standard procedure, which begins with the individual identification of each chromosome, followed by a point and a number (e.g., MQTL1A.1, MQTL1A.2).

The physical positions were obtained for each MQTL using the available sequences of each marker flanking the MQTLs. These nucleotide sequences were retrieved for each marker, using either the published studies or online databases, including GrainGenes (https://wheat.pw.usda.gov/GG3/), Gramene (https://www.gramene.org/), and CerealsDB (https://www.cerealsdb.uk.net/cerealgenomics/CerealsDB/indexNEW.php). Consequently, these nucleotide sequences were subjected to BLASTN searches against the reference genome of wheat available in the Ensemble Plants database (https://plants.ensembl.org/index.html). Genomic regions of some SNPs were obtained from the URGI-JBrowse wheat genome browser (https://wheat-urgi.versailles.inra.fr/).

Furthermore, data from 32 independent GWA studies (involving 10 GWA studies on DS, 4 on HS, 9 on SS, 2 on PHS, and 1 on AS, 6 studies including both DS and HS; no GWAS study was available for WS) were collected to validate the MQTLs discovered in the current study (Table S1). GWA studies, involving durum wheat, hexaploid wheat (e.g., spring and winter wheat), and synthetic hexaploid wheat were conducted across 11 countries. The population size ranged from 70^88^ to 717^89^. The population size, abiotic stress associated, genotyping platform, number of SNP markers employed, and significant markers identified in individual studies are all listed in Table S1. The physical positions of identified significant markers were retrieved from source studies or databases (such as the URGI-JBrowse wheat genome browser and CerealsDB) and compared to the physical coordinates of MQTLs identified on individual chromosomes. MQTLs co-localizing with at least one significant marker were accepted as GWAS validated/verified MQTLs.

### Co-localization of known genes with MQTLs

The association between MQTLs and known genes involved in stress tolerance was also investigated. Sequences of these genes or related markers were BLASTed against the wheat reference genome in the Ensemble Plants database to determine their physical positions. The physical positions of known genes were compared with the physical positions of MQTLs to identify the co-localization of MQTLs with known genes.

### Candidate gene mining, expression analysis, and promoter analysis

MQTLs each conferring tolerance to five or all six abiotic stresses were selected for the identification of underlying gene models. Of the selected MQTLs, MQTLs with a physical CI of ≤ 2 Mb were directly utilized for gene mining. Only a 2 Mb physical region (1 Mb physical region on either side of the MQTL peak) was examined for the availability of gene models for the remaining MQTLs with longer physical CI. The peak genomic positions of the MQTLs were estimated using the formula proposed by our group recently^26^, which is as follows:

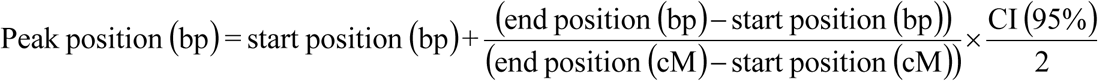

The InterPro database (https://www.ebi.ac.uk/interpro/) was used to obtain descriptions of the functions of discovered gene models. In silico expression analysis of the available gene models was performed using transcriptome data from the wheat expression browser (expression and visualisation platform) powered by expVIP (http://www.wheat-expression.com;^90^). For this purpose, the following relevant expression datasets were utilized: (i) ‘drought and heat stress time-course in seedlings’^91^, (ii) ‘seedlings with PEG to simulate drought’^90^, (iii) ‘spikes with water stress’^90^, and (iv) ‘grain developmental time-course with 4A dormancy QTL’^92^. The first dataset^91^ included differential gene expression in the drought and heat tolerant wheat cultivar TAM107 grown under DS (20% PEG-6000), HS (400C), and both DS and HS conditions (40°C and 20%, PEG-6000), with leaf samples collected separately at 1 and 6 hours after treatment. Seedlings in normal growth conditions (well-watered, 22°C) were used as controls in all of the experiments. The second dataset^90^ consisted of differential expression of genes in two cultivars (Gemmiza10 and Giza168) grown under DS (PEG 6000) with samples from seedlings collected 2 and 12 h after treatment. The third dataset^90^ consisted of differential expression of genes in a doubled haploid population (Westonia x Kauz) grown under water stress conditions with samples collected from the spike at the early booting stage. The fourth dataset^92^ consisted of expression data from 9 NILs (segregating for the 4A dormancy QTL region) and four parental varieties (Baxter, Chara, Westonia, and Yipti) with grain samples collected as 15, 25, and 35 days post-anthesis. The remaining three abiotic stresses, SS, WS, and AL, did not have expression datasets accessible from expVIP. The expression data was given as log2 transformed TPM (transcripts per million) values. Only genes with a substantial fold change (FC ≥ 2 or FC ≤ − 2 TPM values) compared to control were considered differentially expressed. To represent gene expression patterns, heat maps were created using the web tool Morpheus (https://software.broadinstitute.org/morpheus/).

The 1500-bp nucleotide sequences upstream of the translational start site (i.e., ATG) were retrieved from the Ensemble Plants database and uploaded to the PlantCARE database^93^ for cis-regulatory elements (CREs) prediction and potential activities. Only response elements available only on the sense strand with a matrix value of 5 were acceptable, as reported in Sharma et al.^94^.

### Revealing conserved genomic regions in different cereals associated with multiple abiotic stress tolerances

Some of the most promising MQTLs (each conferring tolerance to 5 or all 6 stresses in question) were selected for identification of conserved genomic regions (including ortho- or syntenic MQTL regions) among the wheat, rice, and maize genomes. This analysis involved the following two steps: (i) gene models from the selected MQTL regions were BLASTed against rice and maize genomes at Ensemble Plants to discover conserved genomic regions, and (ii) the matching rice and maize orthologues were retrieved from the database with their genomic coordinates. In order to identify the ortho-MQTLs from the genomic regions conserved among all the three cereals, genomic positions of gene models underlying wheat MQTLs were compared with the orthologous genes available from rice MQTLs for SS tolerance^41^, DS tolerance^42,43^, HS tolerance^44,45^ and maize MQTLs for DS tolerance^38,95,39,40^. MQTLs containing similar gene models located on conserved genomic regions of wheat, barley, rice, and maize were considered ortho-MQTLs.

## Supporting information

E:\The Papers\Meta QTL Paper\MAST MQTL paper final version\Scientific report

## Acknowledgments

Thanks are due to USAID-sponsored Grain Research and Innovation (GRAIN) project for providing GRAIN graduate degree scholarship to MJT for his PhD programme, Department of Science and Technology (DST), New Delhi, India for providing INSPIRE fellowship to DKS and to Head, Department of Plant Breeding and Genetics, Punjab Agricultural University, Ludhiana, (India) for providing necessary facilities.

## Author contributions

AS, MJT and DKS conceived and planned this study. MJT, DKS, KSS, NP, SG and JT performed the literature search, retrieved data, developed a consensus map and conducted the meta-analysis. DKS helped MJT in the analysis and interpretation of results and writing of the first draft of the manuscript. All authors have read and agreed to the final version of the manuscript.

## Data availability statement

Data generated or analysed during this study are included in this published article (and its Supplementary Information files). Datasets are also available from the MJT (Email ID) and DKS (Email ID) on reasonable request.

## Ethics declarations

Not applicable.

## Consent to participate

Not applicable.

## Additional information

**Table 1.** MQTLs providing tolerance to five or all six stresses under study

| **MQTL** | **From (cM)** | **Left marker** | **Right marker** | **Number of QTLs (number of studies)** | **Avg. PVE** | **Associated stress** |
| --- | --- | --- | --- | --- | --- | --- |
| MQTL1B.3 | 65.69-65.96 | BS00082577_51 | RFL_Contig5937_1699 | 37 (17) | 7.89 | AS (2), D+H (7), DS (13), HS (3), PHS (2), SS (10) |
| <sup>a</sup>MQTL1D.4 | 90.8-91.96 | BS00110144_51 | AX-94558596 | 23 (11) | 16.39 | AS (7), D+H (2), DS (1), HS (5), SS (6), WS (3) |
| MQTL2A.5 | 157.81-157.89 | wsnp_Ex_c770_1514612 | Excalibur_c3084_1843 | 44 (14) | 10.7 | AS (1), D+H (1), HS (27), SS (10), WS (5) |
| MQTL2B.3 | 63.78-63.98 | BobWhite_c31708_99 | Excalibur_c4349_1050 | 84 (34) | 10.12 | AS (5), D+H (18), DS (10), HS (22), PHS (4), SS (20), WS (4) |
| MQTL2B.4 | 66.49-67.8 | Kukri_c34861_523 | Kukri_c5497_312 | 13 (12) | 8.01 | AS (3), D+H (4), DS (2), HS (1), PHS (1), SS (2) |
| MQTL2B.6 | 79.65-80.15 | RAC875_c98387_130 | Kukri_rep_c109981_150 | 16 (9) | 11.76 | D+H (2), DS (2), HS (1), PHS (1), SS (5), WS (5) |
| MQTL2D.4 | 61.3-61.38 | Xwmc453.1 | Xcfd56 | 33 (14) | 8.89 | AS (1), D+H (1), DS (10), HS (6), PHS (1), SS (14) |
| <sup>a</sup>MQTL2D.5 | 77.81-78.25 | Excalibur_rep_c107558_367 | GENE-0875_887 | 25 (12) | 21.66 | AS (4), D+H (3), DS (3), HS (4), PHS (1), SS (10) |
| <sup>a</sup>MQTL3A.1 | 180.39-180.4 | BobWhite_c2448_96 | Excalibur_c47078_512 | 70 (35) | 12.19 | AS (6), D+H (9), DS (8), HS (10), PHS (13), SS (21), WS (3) |
| MQTL3B.3 | 103.68-106.27 | AX-94385276 | Tdurum_contig80344_144 | 15 (8) | 7.72 | AS (1), DS (1), HS (3), SS (9), WS (1) |
| MQTL3B.4 | 117.12-117.21 | AX-95131037 | BS00021992_51 | 107 (25) | 8.78 | AS (3), D+H (17), DS (16), HS (23), PHS (6), SS (37), WS (5) |
| MQTL3B.5 | 124.66-125.5 | AX-95020422 | wsnp_Ra_c16264_24873670 | 18 (7) | 9.03 | AS (1), D+H (4), HS (2), SS (10), WS (1) |
| MQTL3B.6 | 155.98-156.42 | Ku_c30806_235 | Xksug53 | 9 (6) | 10.57 | AS (4), D+H (1), HS (2), PHS (1), SS (1) |
| MQTL3B.7 | 202.00-202.33 | AX-94458019 | BS00048754_51 | 16 (9) | 11.78 | D+H (6), HS (1), PHS (2), SS (6), WS (1) |
| <sup>a</sup>MQTL4A.2 | 115.90-116.03 | Tdurum_contig1868_249 | wsnp_BE405275A-Ta_1_1 | 79 (32) | 12.95 | AL (9), D+H (10), DS (18), HS (9), SS (18), WS (1) |
| MQTL4A.5 | 154.63-154.63 | BS00040648_51 | RAC875_c1022_3059 | 16 (9) | 9.11 | AL (2), D+H (2), PHS (4), SS (8) |
| MQTL5A.2 | 117.42-118.07 | BS00100510_51 | Tdurum_contig 29286_319 | 69 (24) | 10.11 | AS (3), D+H (9), DS (26), HS (10), PHS (2), SS (14), WS (5) |
| MQTL5B.5 | 121.79-122.3 | Excalibur_rep_c104627_106 | AX-95113708 | 54 (25) | 9.25 | D+H (11), DS (4), HS (14), PHS (1), SS (20), WS (4) |
| MQTL6B.3 | 92.89-93.49 | BS00088277_51 | AX-94404945 | 51 (22) | 8.47 | AS (4), D+H (1), DS (23), HS (6), PHS (1), SS (16) |
| MQTL7A.4 | 107.20-107.65 | AX-95205723 | AX-94402237 | 27 (14) | 11.34 | AS (5), D+H (1), DS (10), HS (3), PHS (1), SS (7) |
| MQTL7A.5 | 122.91-123.40 | AX-95260715 | AX-94457603 | 47 (22) | 10.52 | AS (3), D+H (8), DS (21), HS (6), PHS (1), SS (8) |
| <sup>a</sup>MQTL7A.6 | 127.9-128.38 | AX-94917420 | AX-95206892 | 16 (11) | 15.24 | AS (2), D+H (4), DS (3), HS (1), PHS (1), SS (5) |
| MQTL7A.7 | 130.35-130.83 | AX-95226392 | AX-94423924 | 13 (10) | 11.6 | D+H (2), DS (4), HS (2), PHS (1), SS (2), WS (2) |
| MQTL7B.3 | 110.98-111.48 | Tdurum_contig47317_100 | BS00066647_51 | 61 (24) | 9.71 | AS (2), D+H (18), DS (8), HS (16), PHS (1), SS (9), WS (7) |
| MQTL7B.4 | 124.39-125.17 | BS00064367_51 | IAAV8521 | 11 (8) | 12.75 | D+H (1), DS (2), HS (1), PHS (1), SS (5), WS (1) |
| MQTL7D.1 | 32.05-34.75 | Xgwm635 | AX-94527430 | 13 (10) | 11.64 | D+H (1), DS (1), HS (5), PHS (1), SS (4), WS (1) |
| MQTL7D.2 | 70.80-72.57 | D_GA8KES401ANMPM_140 | BobWhite_c5654_231 | 26 (12) | 11.26 | D+H (5), DS (5), HS (4), PHS (6), SS (4), WS (2) |
| MQTL7D.3 | 81.29-81.8 | D_contig12156_209 | Kukri_c20975_765 | 18 (13) | 11.94 | AS (2), D+H (6), DS (1), HS (2), PHS (2), SS (5) |

<sup>a</sup>Promising MQTLs selected for breeding programmes.

**Table 2.** Important differentially expressed candidate genes detected in the current study

| MQTL | Candidate genes | Gene start (bp) | Gene end (bp) | GO term name | Function description |
| --- | --- | --- | --- | --- | --- |
| MQTL2B.6 | TraesCS2B02G319600 | 455586854 | 455589182 | integral component of membrane | Late embryogenesis abundant protein |
| MQTL2D.5 | TraesCS2D02G211100 | 168234746 | 168236329 | integral component of membrane | Major facilitator, sugar transporter-like |
| MQTL3B.3 | TraesCS3B02G029900 | 13744964 | 13749320 | diacylglycerol O-acyltransferase activity | O-acyltransferase, WSD1 |
| MQTL3B.4 | TraesCS3B02G039900 | 19263438 | 19270330 | - | Transmembrane protein |
| MQTL3B.5 | TraesCS3B02G040300 | 19869669 | 19872504 | - | Protein of unknown function DUF1666 |
| MQTL3B.7 | TraesCS3B02G466000 | 708516550 | 708520943 | integral component of membrane | Protein of unknown function DUF247 |
| MQTL4A.2 | TraesCS4A02G017700 | 11716257 | 11718716 | nucleus | Zinc finger protein |
| MQTL5A.2 | TraesCS5A02G404900 | 596552839 | 596553578 | - | Uncharacterised conserved protein UCP031279 |
| MQTL7A.4 | TraesCS7A02G143900 | 94786762 | 94791447 | plasma membrane | - |
| MQTL7B.4 | TraesCS7B02G408900 | 678124719 | 678128411 | integral component of membrane | Fatty acid hydroxylase |
| MQTL7D.1 | TraesCS7D02G079400 | 46838388 | 46843665 | ubiquitin-protein transferase activity | Protein kinase domain |

